# The antiparasitics ivermectin and moxidectin trigger genomic and transcriptomic adaptation in bacteria

**DOI:** 10.64898/2026.02.02.703215

**Authors:** Julian Dommann, Georgios Kokotos, Daniel Ballmer, Christian Beisel, Jennifer Keiser, Pierre H. H. Schneeberger

## Abstract

The macrocyclic lactone anthelmintics ivermectin and moxidectin are widely used for parasite control and show antibacterial activity *in vitro*. Given their structural relatedness to macrolide antibiotics, their broad use raises concerns about macrolide cross-resistance and altered bacterial physiology, yet the impact on gut bacterial isolates and underlying mechanisms remains poorly defined. Here, we combined Oxford Nanopore whole genome sequencing with Illumina RNA sequencing across six bacterial isolates analyzed as unexposed controls and as derivatives repeatedly exposed to ivermectin or moxidectin to identify genomic and transcriptomic signatures associated with exposure to these drugs. High-quality genome assemblies enabled integrated analyses across species with distinct cellular architectures and antimicrobial resistance gene arsenals. Across the panel, exposure was associated with i) widespread modulation of ribosome-linked genes and pathways, the molecular target of macrolide antibiotics; ii) context-dependent tuning of pre-existing antimicrobial resistance gene arsenals, such as *macB* and *mdlB*; and iii) recurrent alterations in potassium transport systems (*ktrA* and *trkA*). These findings suggest that repeated ivermectin or moxidectin exposure can reshape bacterial physiology in ways that may, in specific genetic backgrounds, co-modulate resistance-associated traits.

## Introduction

Orally administered chemotherapies are cornerstones of both clinical care and large-scale public health programs. While orally administered medicines ideally target a specific range of pathogens or treat a specific condition, they often represent or resemble natural compounds^1,2^. Over the last two decades, about half of approved drugs have been natural products or derivatives, often of microbial origin, and likely evolved as competitive tools in densely populated, contested ecological niches^2-4^. Irrespective of the approved purpose of these drugs or derivatives thereof, many orally administered drugs, especially when of natural origin, may elicit an antimicrobial effect on the human gut bacteria that they inevitably encounter prior to absorption^5,6^. Human gut bacteria represent an indispensable part of human physiology aiding in digestion, pathogen protection and formation of immune responses. The vast taxonomic diversity and abundance, as well as their genomic and functional potential even created the concept of the human body as a meta-organism^7,8^. Disruption of gut homeostasis has been linked to numerous diseases including cancer, cardiovascular and autoimmune diseases^9,10^. Several factors might lead to gut dysbiosis, oral medications, however, represent a primary one^11^. Beyond disrupting gut homeostasis, repeated exposure to orally administered compounds can select for resistance mechanisms in gut bacteria, expanding the gut resistome and the reservoir of transferable resistance determinants, even when the compound is not an antibiotic^12^. Depending on the mechanisms employed, this could furthermore affect the drug’s bioavailability and therefore potentially change efficacy at the individual and population level. Indeed, besides its key role in health, physiology and its impressive pan-genomic potential, the human gut microbiome’s crucial role in first-pass metabolism is still neglected in pre-clinical, clinical testing or during pharmacovigilance of human chemotherapies.

Ivermectin (IV) and moxidectin (MX) were initially veterinary antiparasitics and were later repurposed for human use against parasitic infections. Structurally, the two naturally derived compounds, IV and MX, represent macrocyclic lactones and are therefore related to macrolide antibiotics. Specifically, IV is approved against several parasitic infections including onchocerciasis, lymphatic filariasis, scabies and strongyloidiasis^13,14^. MX is approved against onchocerciasis and in combination with albendazole undergoing clinical development for soil-transmitted helminthiasis^15-17^. The cumulative burden of these diseases extends far beyond a quarter of the world’s population^18-21^. Since these diseases mostly occur in resource-limited settings, the oral chemotherapies are delivered not primarily as individualized treatment but population-wide through mass drug administration (MDA) – and without prior diagnosis^22,23^. These campaigns repeatedly expose whole communities, including healthy individuals, to the same compounds over years, often with limited capacity for monitoring and surveillance^24^. This repeated, population-wide use may create sustained selection pressure on the gut microbiome at a scale rarely seen with other drug classes: even modest antibacterial activity could reshape the community reservoir of antimicrobial resistance genes. Because the gut microbiome is both a major reservoir for resistance determinants and a hub for horizontal gene transfer^25,26^, selection in commensals can translate into resistance risks in opportunistic or enteric pathogens, with disproportionate consequences where access to second-line antibiotics, diagnostics, and follow-up is limited. Moreover, given IV’s potential application in malaria vector control^27,28^, drug microbiome interactions should be systematically characterized. Supporting this rationale, recent studies put forth evidence of antibacterial properties of both IV and MX *in vitro*^*29*^ and further indicate that baseline gut microbiota composition predicts response to albendazole–IV combination therapy for *T. trichiura* and hookworm^30^. At higher absolute IV doses (≥ 15mg), pronounced taxonomic and functional modulation of the human gut microbiome has been reported^31^. Although the collective resistome was largely untouched in the study, individual bacterial species might still employ resistance mechanisms. Indeed, degradation of IV has been shown to occur in *Escherichia coli* (*E. coli*)^32^.

In this study we define potential mechanisms by which IV and MX interact with gut relevant bacteria to understand potential microbiome-mediated effects on resistance and treatment efficacy under repeated, large-scale use. To accomplish this, we employed a combination of genomic Oxford Nanopore Technologies (ONT) sequencing and transcriptomic Illumina sequencing of bacterial isolates that were repeatedly exposed to IV and MX.

## Methods and Materials

### Selection and origin of isolates and drugs

An overview of the bacterial isolates used in this study is provided in Supplementary Table 1. We utilized a panel of commercial (2/6), and clinical isolates (4/6) that differ in their origin and sensitivity to macrolide or lincosamide antibiotics. Drug-resistant isolates were included to assess whether macrolide or lincosamide resistance is associated with altered sensitivity to IV or MX. As lincosamides share their molecular target, the bacterial ribosome, with macrolides, resistance mechanisms often overlap^33^. The two commercial isolates, *Streptococcus salivarius* (*S. salivarius*) and *Streptococcus parasanguinis* (*S. parasanguinis*), were obtained from the German Collection of Microorganisms and Cell Cultures (DSMZ, https://www.dsmz.de/). 3/6 clinical isolates displayed macrolide (*Streptococcus pneumoniae*, abbreviated *S. pneumoniae*) or lincosamide resistance (*Streptococcus mitis* abbreviated *S. mitis; Streptococcus dysgalactiae* abbreviated *S. dysgalactiae*) and were obtained via the Institute for Infectious Diseases (IFIK, Berne, Switzerland). *S. pneumoniae* is primarily a pulmonary pathogen but is also found throughout the upper and lower gastrointestinal tract and is among the six leading pathogens causing lethal lower respiratory infections^34^. The remaining clinical isolate (*E. coli*) originated from a stool sample collected during a clinical trial evaluating the efficacy and safety of MX–albendazole compared to albendazole and IV– albendazole against *T. trichiura* in Côte d’Ivoire in 2021^35^ (NCT04726969; https://clinicaltrials.gov/study/NCT04726969). Stool samples were collected pre-treatment preserved in 20% (v/v) glycerol, shipped to Switzerland on dry ice, and stored at -80 °C until further use. IV (I8898) and MX (PHR1827) were purchased from Merck & Cie (Buchs, Switzerland).

### Bacterial anthelmintic challenging

The detailed procedure to generate the anthelmintic challenged isolates was published elsewhere^29^. Briefly, six isolates (*S. salivarius, S. parasanguinis, S. pneumoniae, S. mitis, S. dysgalactiae*, and *E. coli*) were started from glycerol stocks by inoculating 10 μl into 10 ml liquid medium (BHI with 5% yeast extract) and incubating for 24 h at 37 °C in 5% CO_2_. Over the next days, cultures were passaged in fresh medium containing increasing concentrations of IV or MX (1 μM, 5 μM, 10 μM). At each concentration, cultures were incubated for 24 h and extended to 48 h if they did not reach OD_600_ > 0.8. If OD_600_ > 0.8 was still not reached, the same condition was repeated, up to a total of three passages; if cultures still did not saturate, challenging for that condition was stopped. Isolates that reached OD600 > 0.8 progressed to the next concentration. The final concentration used for both drugs was 10 μM, yielding challenged isolates labelled with the suffix “IVc” (IV challenged) or “MXc” (MX challenged). Bacterial identification after the final challenging round was performed by ONT 16S amplicon sequencing following a previously published procedure^36^.

### Bacterial sample collection for sequencing

All isolates were handled in an anaerobic chamber (Coy Laboratory Products, Michigan, United States) with a gas mixture of 85% N_2_, 10% CO_2_, and 5% H_2_. BHI + 5% yeast extract was used exclusively to cultivate all isolates from glycerol stocks. Stocks were stored at -80 °C. For cultivation, glycerol stocks were thawed on ice, and 10 μl was inoculated into 10ml of medium. For challenged isolates, growth media were supplemented with 10 μM of the corresponding anthelminthic - IV or MX - dissolved in dimethylsulfoxide (DMSO; Sigma-Aldrich, 41640). Saturated cultures were used as inocula for drug incubation assays. Each of the 6 isolates and their two challenged variants (−IVc, −MXc) were inoculated in duplicate at a 1:2000 dilution into: (1) 10 ml BHI + 5% yeast; (2) 10 ml BHI + 5% yeast with 10 μM IV (0.2% DMSO); (3) 10 ml BHI + 5% yeast with 10 μM MX (0.2% DMSO); or (4) 10 ml BHI + 5% yeast with 0.2% DMSO. For each inoculum, 200 μl was transferred to a 96-well optical plate. The optical plate was incubated in a Hidex Sense plate reader (Hidex Oy, Turku, Finland), recording optical density at λ = 600 nm (OD_600_) every 30 min. In parallel, tubes were incubated anaerobically at 37 °C. When a culture reached OD_600_ = 0.5, one of the tubes was removed, split into five aliquots, and pelleted by centrifugation (3000 x *g*, 10 min). Supernatants were discarded, and pellets were snap-frozen in liquid nitrogen, transferred on dry ice, and stored at -80 °C.

### DNA extraction of bacterial isolates

DNA was extracted from frozen bacterial pellets aliquots using the DNeasy PowerSoil Pro Kit (QIAGEN, 47016) according to the manufacturer’s instructions. The entire pellet, originating from ∼2 ml of culture at OD_600_ = 0.5, was used as input. The final elution volume was set to 60 μl. DNA concentration was determined using a Qubit 4 fluorometer (Thermo Fisher Scientific, Q33238) with the 1x dsDNA HS Assay Kit (Thermo Fisher Scientific, Q33231). Eluates were stored at -20 °C until further use.

### RNA extraction of bacterial isolates

RNA was extracted from frozen bacterial pellets using the Monarch Total RNA Miniprep Kit (New England Biolabs, T2010S) according to the manufacturer’s instructions. Two pellet aliquots, originating from ∼4 ml of culture at OD_600_ = 0.5, were used per sample to ensure sufficient RNA yield. The protocol for “Tough-to-Lyse Samples” was followed, employing mechanical lysis in 400 μl 1x DNA/RNA Protection Reagent. The final elution volume was set to 60 μl. RNA concentration was measured using a Qubit 4 fluorometer (Thermo Fisher Scientific, Q33238) with the RNA HS Assay Kit (Thermo Fisher Scientific, Q32852). Purity was assessed via A_260_/A_280_ and A_260_/A_230_ ratios using a NanoDrop One spectrophotometer (Thermo Fisher Scientific, ND-ONE-W). Eluates were stored at -80 °C until further use.

### Nanopore DNA library preparation and sequencing

Nanopore libraries were prepared using the ONT Native Barcoding Kit 96 (SQK-NBD114-96) according to the manufacturer’s instructions. Sequencing was performed in two parallel runs on an ONT PromethION 2 Solo instrument equipped with two PromethION flow cells (FLO-PRO114M, R10.4.1) for 72 h. We targeted 1 Gb of sequenced bases per sample (Phred score > 10), resulting in an expected genome coverage of approximately 200x - 500x.

### Nanopore DNA read processing

Basecalling of the POD5 files was performed using Dorado^37^ (v. 0.7.2) equipped with the model dna_r10.4.1_e8.2_400bps_sup@v4.3.0. We utilized Filtlong^38^ (v. 0.2.1) to filter short reads (< 300 bp). Subsequently, adapters were removed using Dorado’s function “-trim”. Then, we assembled bacterial genomes using Trycycler^39^ (v. 0.5.4) according to the authors’ instructions. We next assessed the completeness and contamination of assembled genomes using CheckM2^40^ (v. 1.2.2). Subsequently, we annotated the genomes using Prokka^41^ (v. 1.14.6) equipped with the TIGRFAMs^42^ (v. 15.0) and Pfam^43^ (v. 35.0) databases. Average Nucleotide Identities (ANI) between the genomes of the same species were calculated using fastANI^44^. Next, MUMmer^45^ (v. 4.0.0) was used to generate whole-genome alignments and perform SNP calling via the “nucmer”, “delta-filter -1” and “show-snps -T” commands. Using all2vcf^46^ (v. 0.7.8), we generated pseudo-VCF files from the MUMmer output (.snps files) using the “mummer” subcommand and the flag “--snps”. We further performed variant calling via minimap2^47^ (v. 2.26) and clair3^48^ (v. 1.0.6) using the reads of each isolate and its own genome as the reference. VCF files of called variants, as well as called SNPs were annotated using eggNOG^49^, SnpEff and SnpSift^50^ (v. 5.2c). The databases for SnpEff was created with the annotated GeneBank genome files from Prokka (“java -jar snpEff.jar build” with the flags “-genbank -v -noCheckCds –noCheckProtein”). All VCF files were subjected to filtering using bcftools with the flags “view -f PASS -i ‘QUAL>=20 && DP>=40 && (TYPE=“snp” || TYPE=“indel”)’”. To plot the sequencing quality, depth, genome coverage and gene content we utilized R-Studio (R-base v. 4.4.3) equipped with dplyr^51^ (v. 1.1.4), ggplot2^52^ (v. 3.5.2), cowplot^53^ (v. 1.1.3) and ggVenndiagram^54^ (v. 1.5.4).

### Illumina RNA library preparation and sequencing

Illumina RNA library preparation and sequencing were performed at the Genomics Facility of the Department of Biosystems Science and Engineering (D-BSSE) in Basel, Switzerland. RNA eluates were adjusted to a concentration of 15-30ng/μl in nuclease-free water (Ultrapure™ Distilled water, Invitrogen, USA) and integrity and quality of the samples was measured via a bioanalyzer (Agilent TapeStation Systems 4200) prior to library preparation. Libraries were prepared using the Stranded Total RNA Prep with Ribo-Zero Plus Microbiome reagents (Illumina, 20072063). Sequencing was performed on an S4 flow cell using an Illumina NovaSeq 6000 instrument targeting 40M paired-end reads per sample (2 x 100 bp).

### Illumina RNA read processing

Raw Illumina reads were processed using Trimmomatic^55^ (v. 0. 39) and the flags “PE -phred33 ILLUMINACLIP:TruSeq3-PE-2.fa:2:30:10 LEADING:3 TRAILING:3 SLIDINGWINDOW:4:15 MINLEN:36”. We subsequently mapped trimmed reads to the constructed genomes using minimap2^47^ (v. 2. 26) with the flags “-ax sr --secondary=no”. Next, we utilized picard^56^ (v. 3.4.0) to deduplicate the alignment files and added read groups via samtools^57^ (v. 1.18) “addreplacerg”. To count the number of transcripts for each gene, we used featureCounts^58^ (v. 2.0.6). Using this output, we performed first performed quality control (sequencing quality, sequencing depth, hierarchical clustering, principal component analysis). Prior to hierarchical clustering and principal component analysis, the count data was log-transformed and normalized to counts per million (CPM). Thereafter, followed differential expression analysis of the unnormalized, untransformed count data in R-Studio (R-base v. 4.4.3) equipped with dplyr^51^ (v. 1.1.4), reshape2^59^ (v. 1.4.4) and DESeq2^60^ (v. 1.46.0). We filtered transcripts, retaining those with a cumulative count > 10 and collapsed technical triplicates via the command “collapseReplicates”. Since we hypothesize a shared molecular mechanism in response to IV and MX in the bacterial cell, we employed the model “countData ∼ treated”, where the term “treated” combines exposure to either IV or MX. This agrees with our hypothesis and facilitates estimation of dispersion in DESeq2. The significance threshold for differential expression analysis was adjusted to alpha = 0.1. Lastly, differentially expressed genes were annotated via eggNOG. Using the output from differential expression analysis and the KEGG ortholog (KO) annotations generated via eggNOG, we next performed Gene Set Enrichment Analysis (GSEA) with alpha set to 0.1. GSEA was performed in R equipped with clusterProfiler^61^ (v. 4.14.6). To create illustrations, we utilized ggplot2^52^ (v. 3.5.2), ggrepel^62^ (v. 0.9.6), ggVenndiagram^54^ (v. 1.5.4), pheatmap^63^ (v. 1.0.13) and cowplot^53^ (v. 1.1.3).

## Results

### Single-nucleotide polymorphisms (SNPs) in response to anthelmintic exposure

Nanopore sequencing of 18 bacterial isolates – encompassing 6 bacterial species and 12 derived isolates that were challenged with either IV or MX – yielded reads with a median quality score > 14 and sequencing depth ranging between 1.1 – 3.5 Gb per isolate (Supplementary Figure 1). This resulted in high-quality assemblies characterized by genome coverage of > 200x and completeness > 99.98 and contamination < 0.26 across sequenced isolates (Supplementary Figure 2, Supplementary Table 2). Comparing the average nucleotide identity (ANI) between the original isolate subjected to DMSO, and the two challenged variants subjected to IV or MX at 10 μM, we found ANIs for all species ranging from 99.9929% to 99.9999% (Supplementary Table 3). Furthermore, gene content of the original and challenged isolates does not appear to be different with a maximum of two genes being unique to an isolate (Supplementary Figure 3). Additionally, transcriptomic read mapping to these genes showed reads in all derivatives, indicating that the unique genes (*glnL, glnG, glnR, glyG, thrC, secA* and two hypothetical) are likely due to assembly artefacts rather than true gene absence. Subsequently, we investigated single nucleotide polymorphisms (SNPs) occurring in the challenged genomes in respect to the unchallenged genome. The full dataset including all detected SNPs and genomic variants is available in Supplementary File S1. In response to IV exposure, we identified 336 SNPs across the 6 species. 32/336 SNPs occurred on coding regions and they were classified as missense (MS), nonsense (NS), frameshift (FS) or loss of stop codon (SL) SNPs (Table 1, Figure 1). 1/32 occurred in *E. coli*, 11/32 in *S. dysgalactiae*, 5/32 in *S. mitis*, 1/32 in *S. parasanguinis*, 8/32 in *S. pneumoniae* and 6/32 in *S. salivarius*. Across species, SNPs mapped to a limited set of COG pathways, dominated by transcription (*copY, hrcA, tagU/cps4A, glnR, sorC, glcR/fcsR*), translation/ribosome biogenesis (*prmC, prmA, rplR, rpsD*), and inorganic ion transport and metabolism (*ykoD, ktrA, trkA*). Additional variants affected replication/recombination/repair (*E. coli mutL* and *S. dysgalactiae ruvB/dnaX*), as well as multiple metabolic pathways including energy production and conversion (*S. dysgalactiae pta, S. mitis mleA/sfcA*), nucleotide transport and metabolism (*S. dysgalactiae pyrC, pyrD*), and amino acid transport and metabolism (*S. dysgalactiae thrC, S. mitis glnA*). Species-specific single hits were also observed in cell division (*S. dysgalactiae divIVA*), defense mechanisms (*S. dysgalactiae femX/murM*), and cell wall/membrane/envelope biogenesis (*S. mitis murA* and *S. salivarius arnC/cpsJ*), along with *S. pneumoniae* variants in coenzyme metabolism (*fpgS/folC*), secondary metabolite related functions (*axeA/cah*), and intracellular trafficking/secretion (*secA*). Notably, *secA* also showed a highly frequent genomic variant representing the reversed genotype of the detected SNP (allele frequency = 0.914), which was the only such high frequency variant observed across all species. In response to MX exposure, we identified 114 SNPs across the 6 isolates, of which 44 were found to cause mutations on coding regions (Table 2, Figure 1). 1/44 occurred in *E. coli*, 14/44 in *S. dysgalactiae*, 8/44 in *S. mitis*, 9/44 in *S. parasanguinis*, 6/44 in *S. pneumoniae* and 6/44 in *S. salivarius*. Across species, SNPs mapped mainly to five recurring COG pathways, dominated by inorganic ion transport and metabolism (*ykoD, ktrA, trkA, btuD/rgpD, nrgA/amt*), carbohydrate transport and metabolism (*glgX/pulA, agaS, manX/manL, yutF/nagD, lacZ/lacL*), transcription (*copY, nrdR, relA, ctsR, glnR*), translation and ribosomal structure/biogenesis (*prmC, rsmF, prmA, rnj2/rnjB*), and defense mechanisms (*msbA, rny, yheH/mdlB*). Additional variants affected replication, recombination and repair (*S. dysgalactiae nth* and *ruvB/dnaX*) and multiple metabolic pathways, including energy production and conversion (*S. dysgalactiae pta, S. mitis mleA/sfcA, S. salivarius pflB*), nucleotide transport and metabolism (*S. dysgalactiae pyrD* and *S. parasanguinis hpt*), and amino acid transport and metabolism (*S. dysgalactiae dapE* and *thrC, S. parasanguinis ilvA*). Species-specific single hits were also observed in cell division (*S. dysgalactiae divIVA*), coenzyme transport and metabolism (*S. pneumoniae fpgS/folC*), cell wall/membrane/envelope biogenesis (*S. pneumoniae ywqC/cps4C*), posttranslational modification/protein turnover/chaperones (*S. salivarius ureD*), and lipid transport and metabolism / secondary metabolite related functions (*S. salivarius fabG*), alongside several loci that were unknown or unassigned in this dataset (*S. mitis cmpC, S. parasanguinis xylG/mglA, E. coli norR* and *S. dysgalactiae skc*).

**Table 1.**
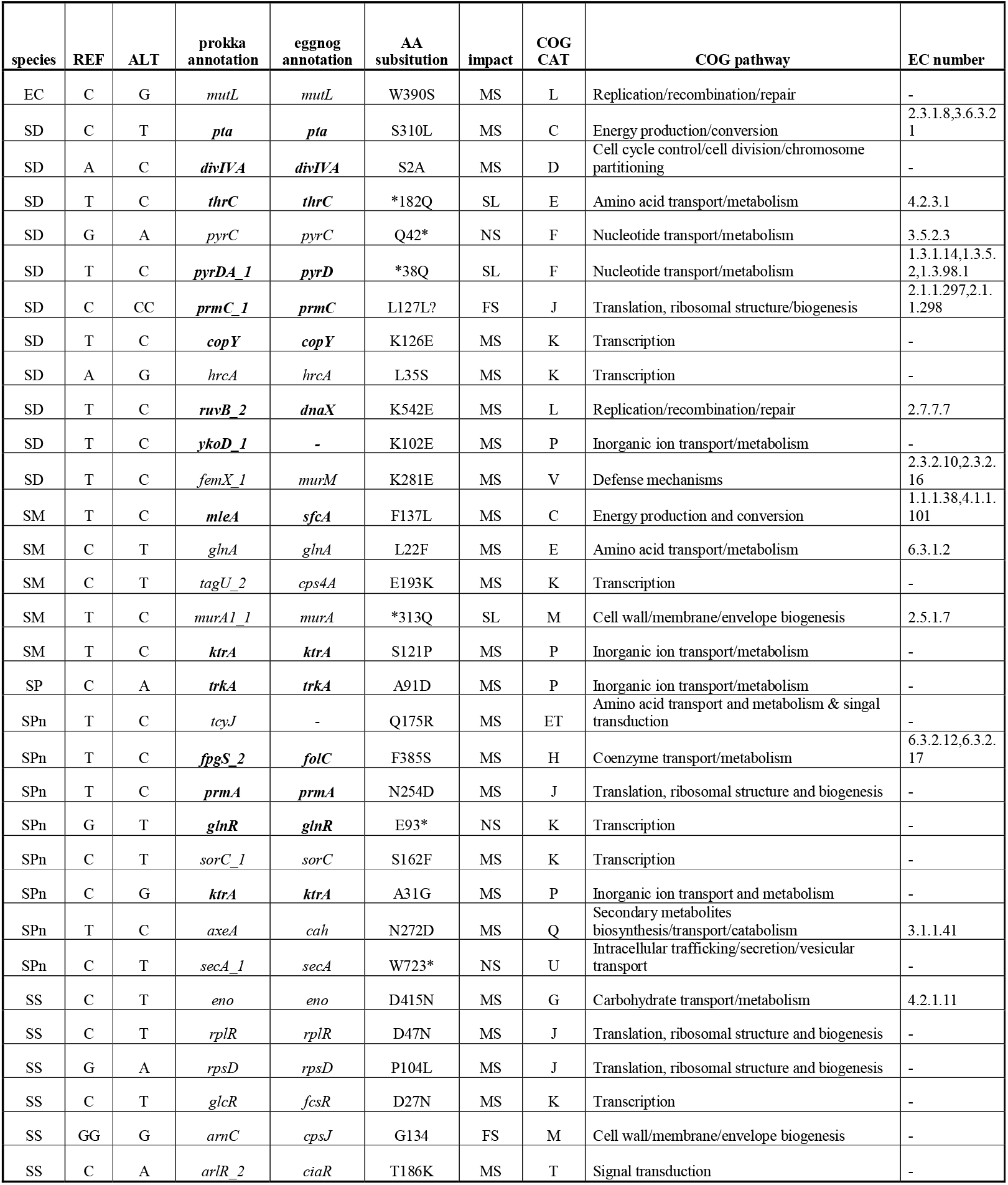
Overview of 32 detected single-nucleotide polymorphisms (SNPs) in six bacterial isolates after prolonged exposure to ivermectin. Genes printed in bold were also found mutated after moxidectin exposure (Table 3). EC = *Escherichia coli*, SD = *Streptococcus dysgalactiae*, SM = *Streptococcus mitis*, SP = *Streptococcus parasanguinis*, SPn = *Streptococcus pneumoniae*, SS = *Streptococcus salivarius*. REF = reference nucleotide (control genome), ALT = alternative nucleotide (exposed genome), AA = amino acid, COG CAT = clusters of orthologous genes category, EC number = enzyme commission number. MS = missense, NS = nonsense, FS = frame shift, SL = stop lost.

**Table 2.**
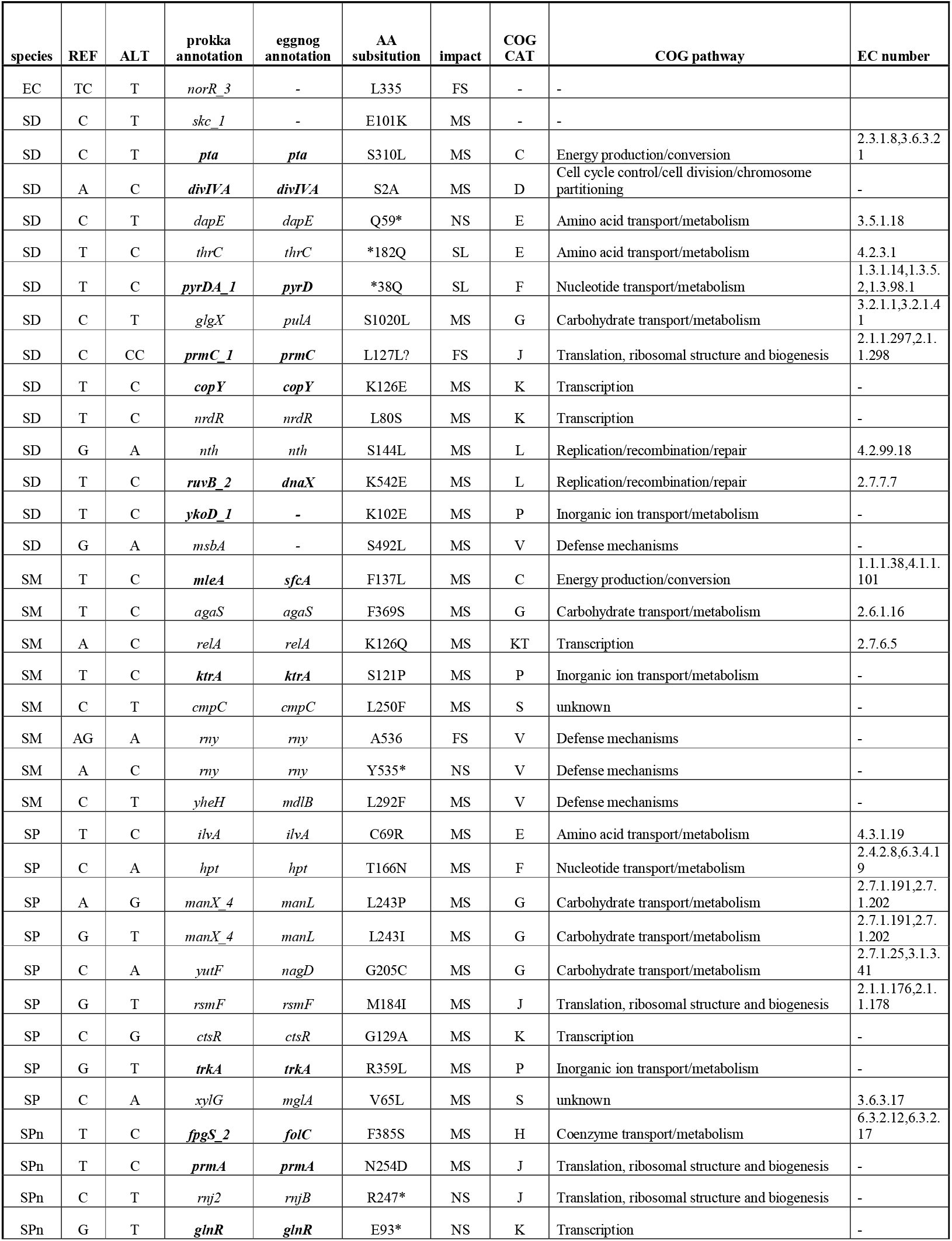
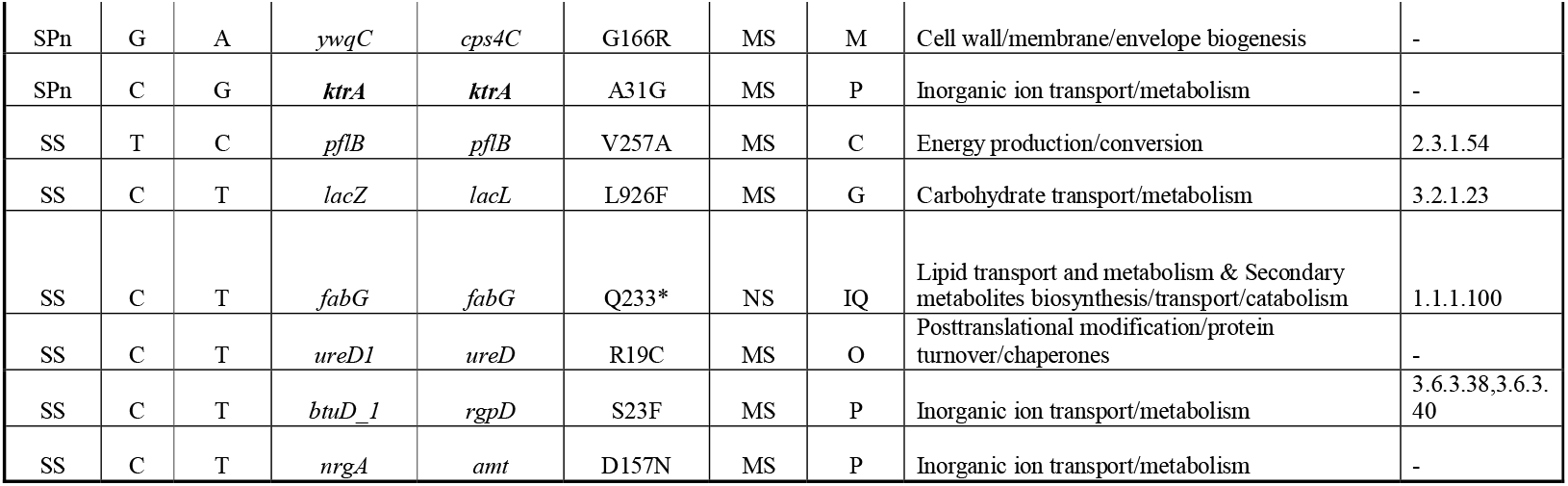
Overview of 44 detected single-nucleotide polymorphisms (SNPs) in six bacterial isolates after prolonged exposure to moxidectin. Genes printed in bold were also found mutated after ivermectin exposure (Table 2). EC = *Escherichia coli*, SD = *Streptococcus dysgalactiae*, SM = *Streptococcus mitis*, SP = *Streptococcus parasanguinis*, SPn = *Streptococcus pneumoniae*, SS = *Streptococcus salivarius*. REF = reference nucleotide (control genome), ALT = alternative nucleotide (exposed genome), AA = amino acid, COG CAT = clusters of orthologous genes category, EC number = enzyme commission number. MS = missense, NS = nonsense, FS = frame shift, SL = stop lost.

**Figure 1:**
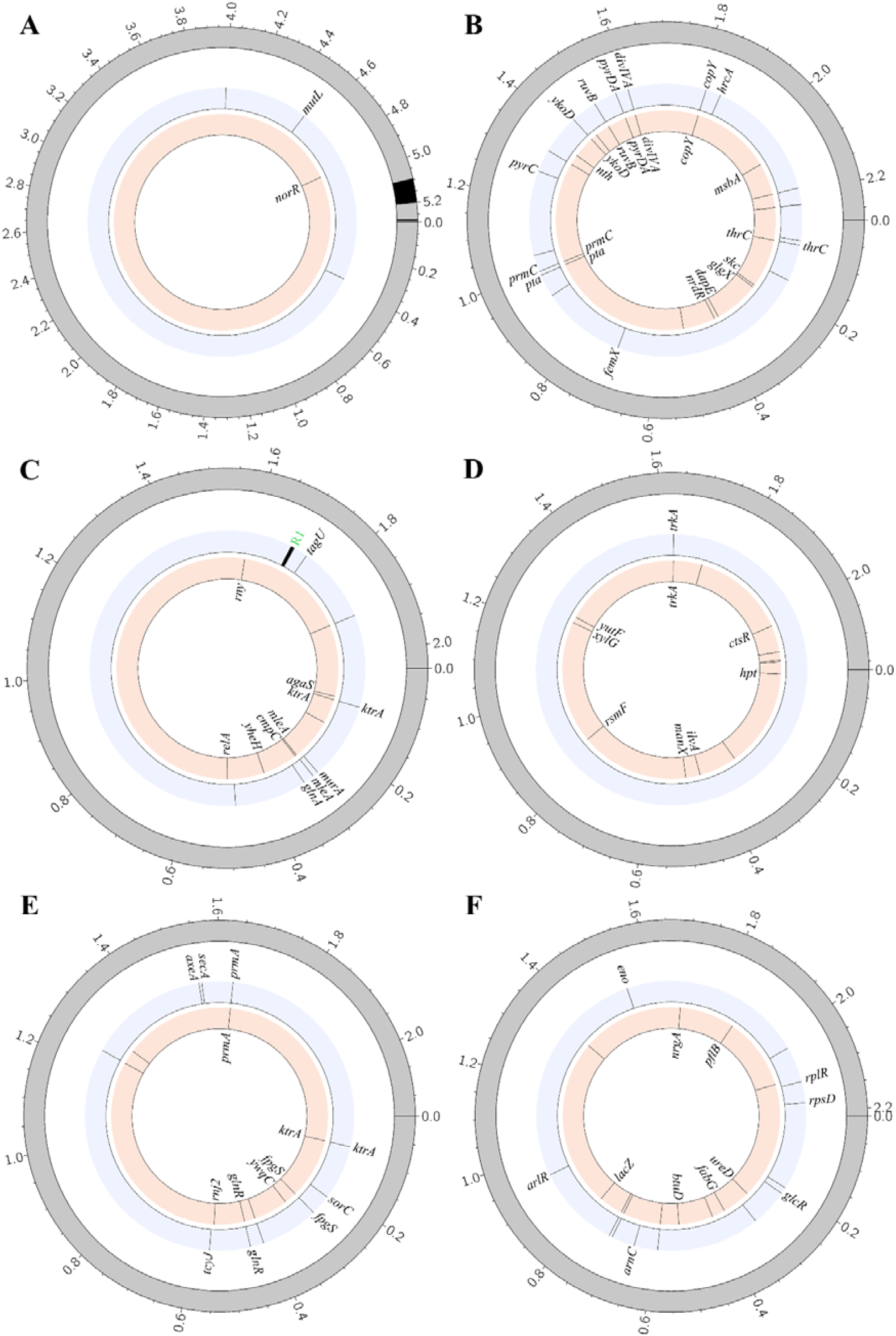
Circularized genomes of the six bacterial species. The outer ring represents the contig(s), colored in grey and black, originating from the assembled control genome (exposed to DMSO only) and genome positions are indicated in megabases (Mb). The inner rings represent the circularized genomes of the anthelmintic-exposed isolates (blue = IVc, orange = MXc). Black lines in the inner rings represent the position of single nucleotide polymorphisms (SNPs) that were detected on that genome respective to the control genome. (**A**) *Escherichia coli*. (**B**) *Streptococcus dysgalactiae*. (**C**) *Streptococcus mitis*. The label R1 (green) refers to an accumulation of SNPs that are localized to one gene coding for a hypothetical protein (HP). (**D**) *Streptococcus parasanguinis*. (**E**) *Streptococcus pneumoniae*. (**F**) *Streptococcus salivarius*.

### Shared and unique gene expression patterns upon anthelmintic exposure

Building on the genomic results, we performed transcriptome analyses to provide functional context and to test whether prolonged exposure to IV or MX is associated with inducible expression programs, potentially linked to AMR mechanisms such as drug efflux, that could contribute to tolerance or resistance. Hence, using the RNA sequencing data we generated for *E. coli, S. dysgalactiae, S. mitis, S. parasanguinis, S. pneumoniae, S. salivarius* and their challenged counterparts (-IVc / -MXc), we performed differential expression analysis and subsequent Gene Set Enrichment Analysis (GSEA). RNA sequencing resulted in > 90% of reads quality scores > Q30 across samples (Supplementary Figure 1). Since each sample was sequenced in three technical replicates, we first assessed the similarity of these replicates as a measure of quality control. Indeed, hierarchical clustering, as well as principal component analysis (PCA) revealed consistency between replicates (Supplementary Figures 4-5). Triplicate consistency was quantified as the Euclidean distance to each condition centroid in PC1–PC2 space (88.3–99.5% variance explained; Supplementary Table 4). Across all species and conditions, replicates clustered tightly (mean distance to condition centroid 0.073–0.683). The only exception was one replicate of *E. coli* (-IVc), which showed markedly higher dispersion (mean 3.50). We next performed differential expression analysis in DESeq2 to identify transcripts which expression is associated with anthelminthic exposure. Since we hypothesize one or more shared molecular targets of IV and MX in bacteria, our employed model combines exposure to either IV or MX. The full differential expression results dataset is available in Supplementary File S2. Differential expression analysis revealed both shared and species-specific transcriptional responses to anthelmintic exposure, with the magnitude of response varying substantially across taxa. *S. dysgalactiae* exhibited the most pronounced transcriptional remodeling (142 differentially expressed genes), followed by *E. coli* (67 genes) and *S. parasanguinis* (65 genes), whereas *S. mitis, S. pneumoniae*, and *S. salivarius* showed progressively more constrained responses.

In *E. coli*, differentially expressed genes clustered into coherent metabolic and transport-associated modules. Members of the *gnt* (*gntK, gntU, gntT*) and *nap* (*napA–D, F–H*) operons were consistently downregulated, while genes involved in amino acid and sulfur metabolism were strongly induced, including the *ast* (*astA–E*) and *cys* (*cysA, C, D, H–J, N, P, T, W*) gene families. Notably, the most strongly downregulated transcripts (log_2_FC < −4.0) were *mdtI, mdtJ*, and *ydhC*, which encode components of intrinsic multidrug efflux systems, indicating a broader physiological adaptation. *S. dysgalactiae* displayed extensive transcriptional restructuring across resistance, metabolic, transport, and translational functions. For instance, genes associated with antimicrobial resistance and stress response showed divergent regulation: *erm, msbA*, and the *lnr* operon (*lnrL/M/N; sagG/H/I*) were downregulated, while *yknX* and *yknY*, encoding components of MacB-family macrolide efflux transporters, and *msrA* were upregulated. In parallel, numerous ribosomal genes were differentially expressed, with downregulation of *rplS, rpmA, rpmF, rpmGA, rpmH, rpsD*, and *rpsU* (log_2_FC ≤ - 1.10), and upregulation of *rplM, rpsM, rsmD*, and *rsmG* (log_2_FC ≥ 1.10), indicating extensive remodeling of translational machinery. Genes involved in carbohydrate utilization, particularly members of the *lac* operon (*lacA–D, F, R, X*), were broadly downregulated, whereas purine biosynthesis genes (*purC, F, L–N*) were upregulated, suggesting altered nucleotide metabolism. Several transport-associated genes were differentially regulated, including upregulation of *btuD* (vitamin B12 import) and *ycjP* (maltose transport), and downregulation of *isdF* (iron transport). In contrast, *S. mitis* exhibited a highly focused transcriptional response, with only seven genes differentially expressed. Among these, *macB* (log_2_FC = 25.73) and *murA* (log_2_FC = 3.86) showed strong induction, linking anthelmintic exposure to macrolide efflux and cell wall biosynthesis pathways, respectively. Consistent with patterns observed in *S. dysgalactiae, lacC* and *lacR* were downregulated. *S. parasanguinis* showed differential expression of genes affecting carbohydrate metabolism, translation, and stress response. Members of the *lac* family (*lacC, E–G, R*) were differentially regulated in both directions, while *rplO*, encoding the 23S rRNA-binding protein L15, was upregulated. In addition, genes involved in amino acid biosynthesis (*leuB–D*) were induced, whereas chaperone (*dnaJ/K, groL/S*), desaturase (*desK/R*), and pyrimidine metabolism (*pyrB, pyrR*) genes were downregulated. More limited responses were observed in *S. pneumoniae* and *S. salivarius*. In *S. pneumoniae*, ten genes were differentially expressed, including bidirectional regulation of *lac* genes (*lacC, E–G*), upregulation of *rnj2*, and downregulation of *thrC. S. salivarius* displayed the smallest response, with only two genes affected: strong upregulation of *glyG* (log_2_FC = 26.20) and downregulation of *btuD* (log_2_FC = −2.45).

We next aimed to complement the gene-level changes with a pathway-level view by applying GSEA (Fig. 2BC) to test whether sets of functionally related genes show coordinated expression shifts under prolonged IV or MX exposure. Across species, we found different trends; gene sets with a significant Normalized Enrichment Score (NES) in only one species or multiple, and if in multiple, equally signed NES or opposite. Aminoacyl-tRNA biosynthesis and biosynthesis of phenylalanine, tyrosine and tryptophan were negatively enriched, whereas purine metabolism and the two-component system were positively enriched in multiple species. There were also species-specific gene sets, for example homologous recombination, protein export and sulfur metabolism for *E. coli*, fructose and mannose metabolism for *S. mitis* or galactose metabolism for *S. parasanguinis*. Interestingly, ABC transporters were negatively associated with anthelmintic exposure in *S. mitis*, but positively in *E. coli*. Most striking is the significant association of ribosomal function across 5 out of 6 species (not in *S. mitis*). Directionality in this gene set is not uniform, as it is positively associated in *S. parasanguinis* and *S. pneumoniae* but negatively associated in *E. coli, S. dysgalactiae* and *S. salivarius*. Except for *S. salivarius*, closer inspection of the gene sets further reveals that over 50% of the core ribosomal gene set of over 40 genes is affected – in *E. coli* over 75% (Supplementary Figure 6). The distribution of gene enrichment scores among significantly enriched gene sets (Figure 2C) confirms uniform distribution to either the positive or negative side of the ranked gene list and reveals patterns that might indicate refined, multi-stepped regulation in some gene sets, such as ABC transporters, protein export, purine metabolism, ribosome, sulfur metabolism and the two-component system, given the presence and distribution of multiple peaks in the density curves.

**Figure 2:**
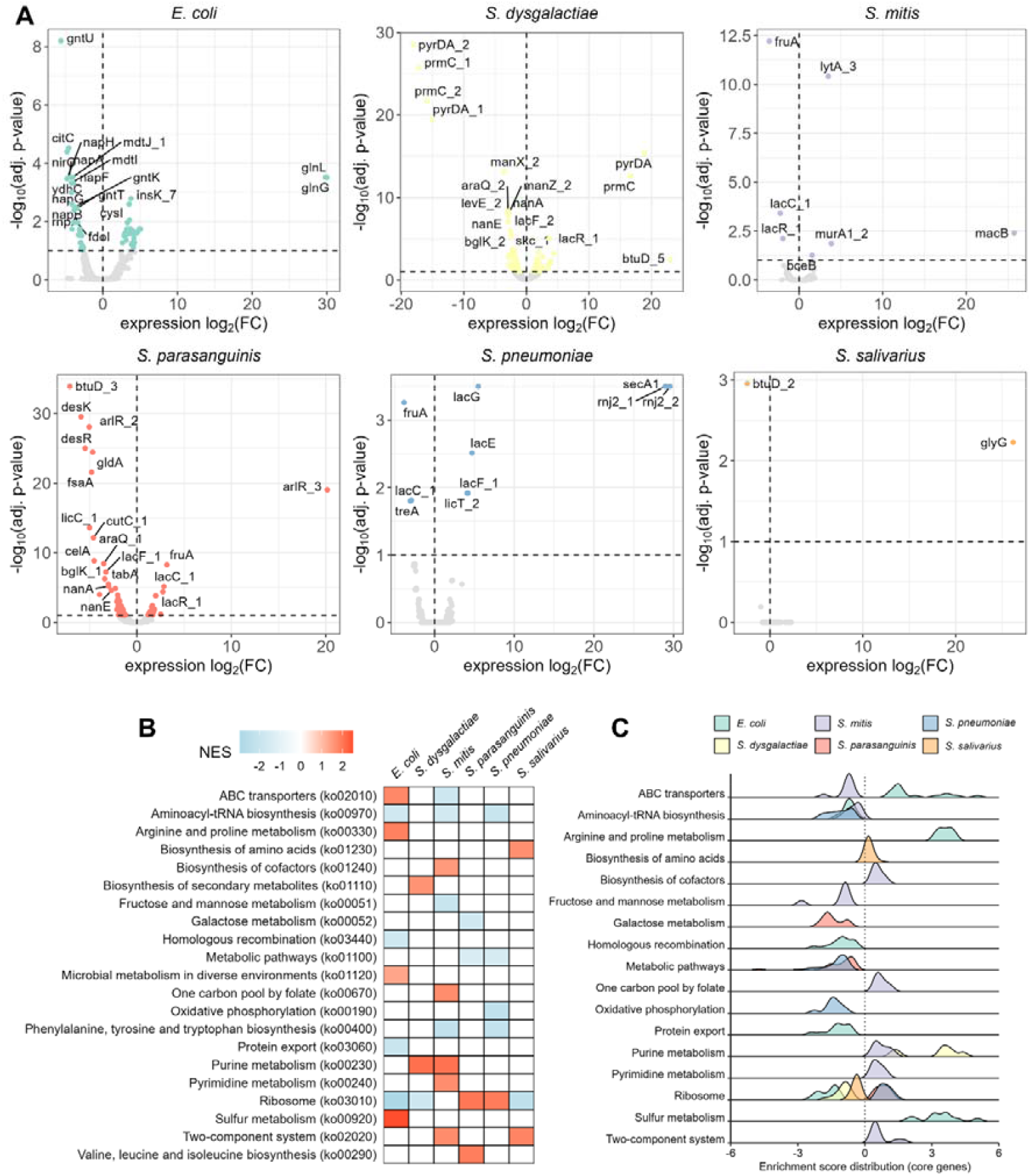
Differential expression patterns of bacterial isolates in response to anthelmintic exposure. (**A**) Volcano plot summarizing the differential expression analysis conducted via DESeq2 by means of log_2_fold-change of transcript expression and statistical significance. Blue coloration indicates a significance threshold of alpha = 0.1 or lower. (**B**) Heatmap of significant Normalized Enrichment Scores (NES) across enriched KEGG ortholog (KO) gene sets. (**C**) Distribution of enrichments scores among gene sets with a significant NES.

## Discussion

IV and MX are widely and frequently used anthelmintics deployed at exceptional scale^64^, repeatedly exposing gut bacterial communities across entire populations. Both have established antibacterial activity *in vitro*^*5,29*^, which may in part relate to their macrocyclic chemical architecture that resembles macrolide antibiotics. Hence, even modest off-target antibacterial activity becomes a public health question: repeated exposure could impose selection pressure on gut bacteria and drive adaptive responses that may i) co-modulate resistance-associated traits, including drug efflux, or ii) alter drug bioavailability, for example through uptake^65^ or drug degradation^32^, as shown for IV in *E*.*coli*^*32*^. To identify genomic and transcriptomic signatures associated with IV or MX exposure, we combined ONT long read genome sequencing with Illumina RNA sequencing across six phylogenetically and phenotypically diverse isolates spanning distinct origins, cellular architectures and antimicrobial resistance arsenals, enabling detection of both shared and species-specific responses.

Across the panel, the dominant patterns were better captured by shared responses than by species-specific ones. However, *E. coli* emerged as the only isolate that was intrinsically insensitive, likely due to limited drug access across the gram-negative cell envelope. We did not observe clear phenotypic or genomic shift after repeated exposure, aligning with previous reports that *E. coli* is insensitive to macrolide antibiotics^66^. Still, a previous study demonstrated degradation of IV by an *E. coli* isolate^32^, implying that at least a fraction of the drug can enter the cell, which is likely associated with the transcriptomic signal observed here, including upregulation of sulfur metabolism (*cys* genes), decreased efflux-associated gene expression (*mdtI, mdtJ, ydhC*), and depletion of the ribosome associated gene set.

Beyond *E. coli*, ribosome-linked alterations emerged as a recurrent pattern across the panel. Multiple independent findings point to a consistent association between exposure to IV or MX and ribosome-linked components including ribosomal gene mutations, differential regulation of ribosomal proteins, and GSEA enrichment of ribosome associated gene sets in five of six species. While this mirrors the functional neighborhood targeted by macrolide and lincosamide antibiotics, these data do not establish a direct molecular interaction with the ribosome; instead, they indicate that ribosome-associated pathways are repeatedly modulated, whether as a direct effect, or a downstream compensatory adjustment. Regardless of direct binding, the consistent convergence on ribosome-linked signatures indicates that these compounds repeatedly perturb the same functional neighborhood that underlies clinically important antibacterial drug responses.

In addition, we observed context-dependent tuning of pre-existing AMR arsenals rather than a uniform directional shift. In some species, AMR-associated genes were downregulated (*erm, msbA, lnr* genes), while selected macrolide efflux related components were upregulated, including *yknY/macB*^67^ in more than one streptococcal species (notably *S. dysgalactiae* and *S. mitis*). This pattern supports a model in which isolates with existing AMR arsenals can reallocate expression toward functions that are beneficial under the specific exposure condition (e.g. macrolide efflux), without implying a single, conserved resistance mechanism or a defined bacterial target for these compounds.

Finally, in isolates where exposure coincided with broader molecular changes, the data were consistent with cellular stress and homeostasis responses. Several gram-positive species, most prominently *S. dysgalactiae* and *S. mitis*, showed higher SNP burdens and stronger transcriptional remodeling, with mutations in genes linked to cellular stress, ribosomal modifications and genome maintenance, including *msbA, prmC*, and *ruvB*^68-70^. A recurrent signal across multiple species was altered potassium transport, with mutations in *ktrA* or *trkA* appearing in independent lineages (including *S. mitis, S. pneumoniae*, and *S. parasanguinis*). This convergence suggests that ion homeostasis and membrane linked physiology are repeatedly affected during prolonged exposure to IV or MX and may contribute to the observed responses. For instance, absence of *ktrA* was previously linked to increased sensitivity to hyperosmotic conditions and susceptibility to aminoglycoside antibiotics and cationic antimicrobials in *S. aureus*^*71*^.

Although phenotypic adaptation of these six isolates under repeated IV or MX exposure was reported previously^29^, this study resolves the associated molecular signatures. Notably, the two species with the strongest transcriptomic remodeling (*E. coli, S. dysgalactiae*), measured by the number of differentially expressed genes, showed the smallest phenotypic shifts, consistent with the idea of compensatory responses. To conclude, our findings can be summarized in four main points: i) *E. coli* appears intrinsically insensitive to IV and MX and therefore shows only limited phenotypic, genomic and transcriptomic responses, consistent with poor drug access across the gram negative-cell envelope; ii) multiple independent lines of evidence point to a reproducible association between exposure to IV or MX and the bacterial ribosome, the molecular target of macrolide and lincosamide antibiotics; iii) in several gram-positive isolates, a pre-existing AMR arsenal appears to be fine-tuned under IV or MX exposure, predominantly through modulation of drug efflux systems and macrolide-associated determinants, such as *macB* or *mdlB*; and iv) ion homeostasis emerges as a recurrently affected function, with independent mutations in *ktrA/trkA* across multiple species linking repeated exposure to altered potassium transport. In the broader context of large-scale IV and MX use, these patterns represent a crucial finding, as they define a limited set of reproducible and testable routes by which off-target exposure could reshape bacterial physiology and, in some settings, co-modulate resistance-associated traits. For drugs used at large scale, selection pressure on the gut microbiome is a relevant consideration that should be critically evaluated.

Our findings define adaptive phenotypic, genomic, and transcriptomic responses to IV and MX under controlled exposure conditions, but do not directly address their population-level or long-term consequences in treated communities. While the convergence of multiple data layers implicates ribosome-associated functions, ion homeostasis, and transport processes, direct causal validation of these interactions was beyond the scope of this study. Future work integrating longitudinal metagenomic analyses with targeted genetic perturbations and direct measurements of compound uptake or transformation will be essential to further resolve these mechanisms.

## Data Availability

The sequencing data (ONT and Illumina shotgun sequencing) generated in this study will be deposited in the NCBI Short Read Archive. Numerical source data underlying all figures, as well as detailed documentation of the bioinformatics analysis can be found under https://github.com/STPHxBioinformatics/IVtranscriptoMX.

## Author Contributions

J.D.: study design, research design, project supervision, experimental work, statistical analyses, figure generation, writing of the initial paper, and paper editing. G.K.: experimental work (bacterial incubations) and paper editing. D.B.: experimental work (ONT DNA sequencing), and paper editing. C.B.: coordination of experimental work (Illumina RNA sequencing), and paper editing. J.K.: study design, research design, project supervision, funding acquisition, and paper editing. P.H.H.S.: study design, research design, project supervision, funding acquisition, and paper editing.

## Acknowledgements

Sequencing library preparation and Illumina sequencing were carried out at the Genomics Facility Basel of the University of Basel and Department of Biosystems Science and Engineering, ETH Zurich. All steps of the computational analysis were performed using the scientific computing centre at the University of Basel (sciCORE; http://scicore.unibas.ch/).

