## Supplementary Materials for "The antiparasitics ivermectin and moxidectin trigger genomic and transcriptomic adaptation in bacteria"

Supplementary Table 1: Overview of the bacterial samples used in this study. Isolate 01 was isolated from a stool sample obtained in the clinical trial registered under NCT04726969. Abbreviations: IFIK = Institute for Infectious Diseases, DSMZ = German Collection of Microorganisms and Cell Cultures.

| **sample** | **species** | **acquisition via** |
| --- | --- | --- |
| 01 | *Escherchia coli* | NCT04726969 |
| 02 | *Streptococcus dysgalactiae* | IFIK, Berne, Switzerland |
| 03 | *Streptococcus mitis* | IFIK, Berne, Switzerland |
| 04 | *Streptococcus parasanguinis* | DMSZ, DSM 6778 |
| 05 | *Streptococcus pneumoniae* | IFIK, Berne, Switzerland |
| 06 | *Streptococcus salivarius* | DMSZ, DSM20067 |

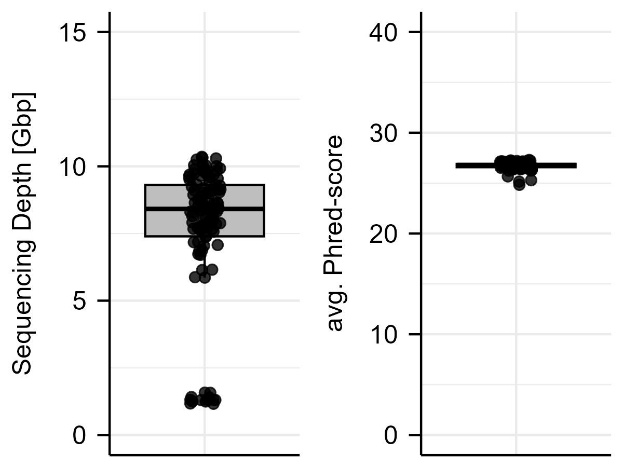

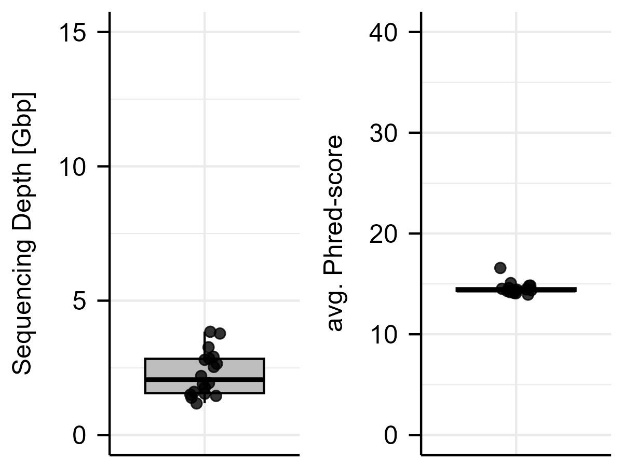
**A**	**B**

Supplementary Figure 1: Sequencing depth and quality, given as Phred-score, of the (A) Oxford Nanopore Technologies genomic and (B) Illumina transcriptomic sequencing runs.

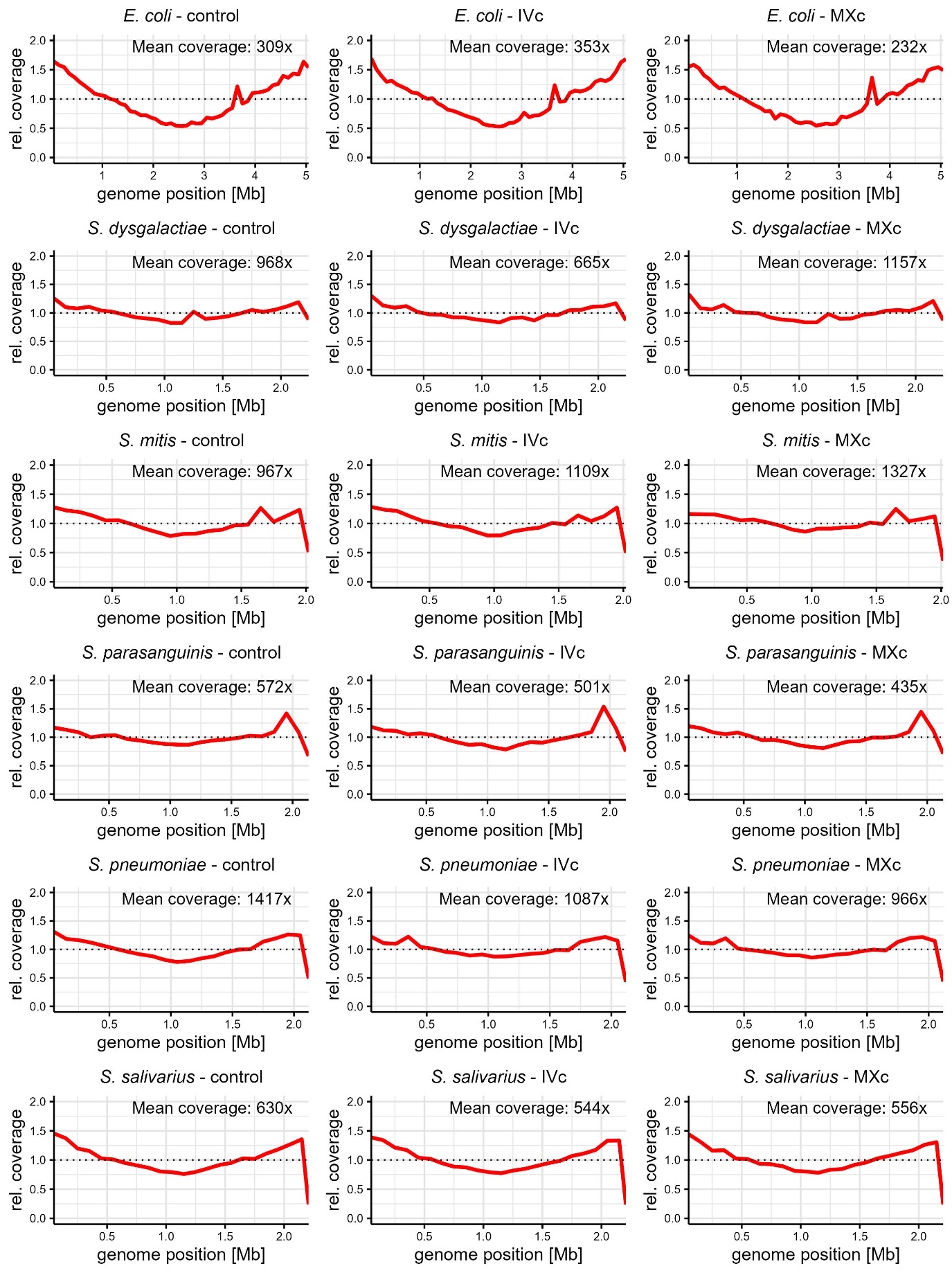

Supplementary Figure 2: Relative genome coverage of 18 bacterial isolates after assembly.

Supplementary Table 2: Overview of assembly quality generated via checkm2.

| **sample** | **completeness [%]** | **contamination [%]** | **genome size [b]** | **no. of contigs** | **contig N50 [b]** | **GC content** |
| --- | --- | --- | --- | --- | --- | --- |
| *E. coli* | 100 | 0 | 5284764 | 5 | 5095383 | 0.51 |
| *E. coli (-IVc)* | 100 | 0 | 5284762 | 5 | 5095381 | 0.51 |
| *E. coli (-MXc)* | 100 | 0 | 5284763 | 5 | 5095382 | 0.51 |
| *S. dysgalactiae* | 100 | 0.26 | 2273984 | 1 | 2273984 | 0.39 |
| *S. dysgalactiae (-IVc)* | 100 | 0.26 | 2273983 | 1 | 2273983 | 0.39 |
| *S. dysgalactiae (-MXc)* | 100 | 0.26 | 2273983 | 1 | 2273983 | 0.39 |
| *S. mitis* | 99.99 | 0.02 | 2039314 | 1 | 2039314 | 0.4 |
| *S. mitis (-IVc)* | 99.99 | 0.02 | 2039080 | 1 | 2039080 | 0.4 |
| *S. mitis (-MXc)* | 99.98 | 0.03 | 2031676 | 1 | 2031676 | 0.4 |
| *S. parasanguinis* | 99.99 | 0.01 | 2164415 | 2 | 2161178 | 0.42 |
| *S. parasanguinis (-IVc)* | 99.99 | 0.01 | 2166517 | 2 | 2163280 | 0.42 |
| *S. parasanguinis (-MXc)* | 99.99 | 0.01 | 2164628 | 2 | 2161391 | 0.42 |
| *S. pneumoniae* | 100 | 0.16 | 2136255 | 1 | 2136255 | 0.4 |
| *S. pneumoniae (-IVc)* | 100 | 0.15 | 2136253 | 1 | 2136253 | 0.4 |
| *S. pneumoniae (-MXc)* | 100 | 0.15 | 2136253 | 1 | 2136253 | 0.4 |
| *S. salivarius* | 99.99 | 0.11 | 2214094 | 1 | 2214094 | 0.4 |
| *S. salivarius (-IVc)* | 99.99 | 0.11 | 2214094 | 1 | 2214094 | 0.4 |
| *S. salivarius (-MXc)* | 99.99 | 0.11 | 2214093 | 1 | 2214093 | 0.4 |

Supplementary Table 3: Average Nucleotide Identities (ANIs) between the control isolates and the anthelmintic challenged counterparts (suffix “-IVc” or “-MXc”). ANIs were calculated via fastani.

| **species** | **versus -IVc [%]** | **versus -MXc [%]** |
| --- | --- | --- |
| *E. coli* | 99.9973 | 99.9999 |
| *S. dysgalactiae* | 99.9978 | 99.9980 |
| *S. mitis* | 99.9993 | 99.9929 |
| *S. parasanguinis* | 99.9991 | 99.9986 |
| *S. pneumoniae* | 99.9986 | 99.9985 |
| *S. salivarius* | 99.9991 | 99.9989 |

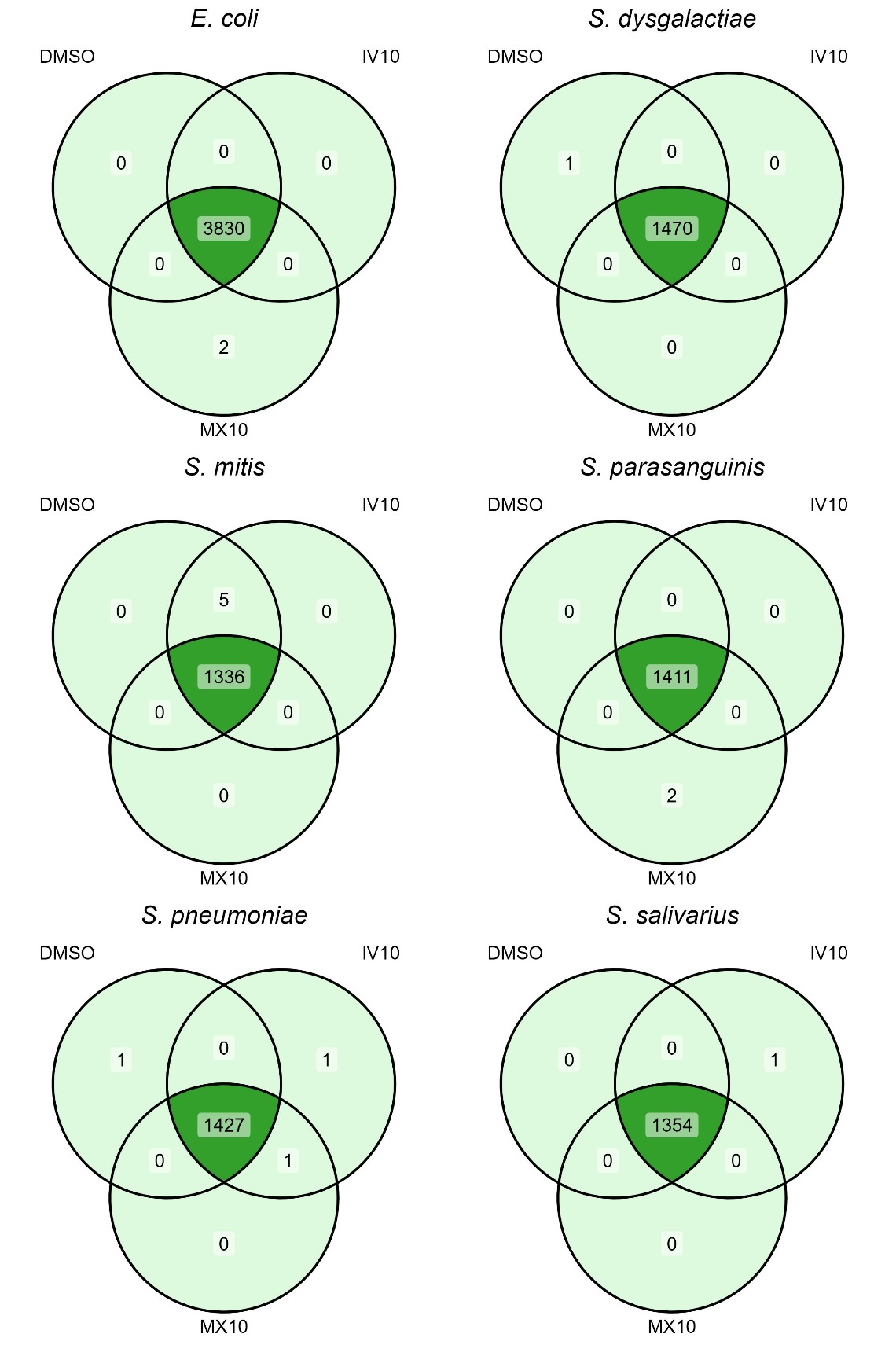

Supplementary Figure 3: Shared and unique genes for unchallenged, IV-challenged and MX-challenged isolates per species, annotated via UniProt, Pfam and TIGRFAMs accessions. Unique genes included glnL (E. coli, MXc), glnG (E. coli), thrC (S. dysgalactiae), two hypothetical proteins (S. parasanguinis, MXc), glnR (S. pneuomoniae), secA (S. pneuomoniae, IVc), and glyG (S. salivarius, IVc).

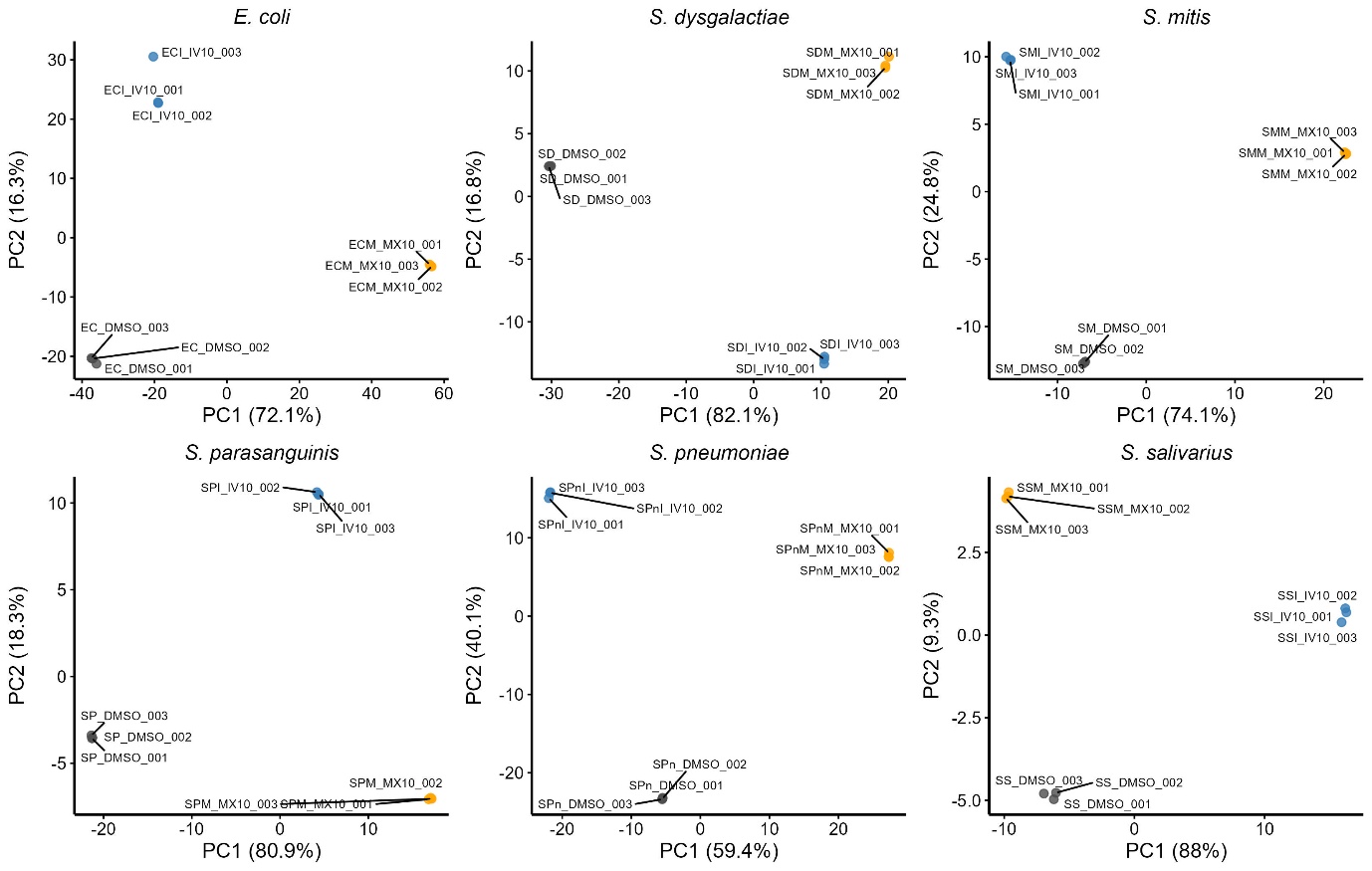

Supplementary Figure 4: PCA ordination of expression profiles of 18 sample triplicates. Abbreviations: PC = principal component, EC = Escherichia coli, SD = Streptococcus dysgalactiae, SM = Streptococcus mitis, SP = Streptococcus parasanguinis, SPn = Streptococcus pneumoniae, SS = Streptococcus salivarius, IV10 = ivermectin at 10µM, MX10 = moxidectin at 10µM.

Supplementary Table 4: Overview of replicate distances in the principal component analysis (PC; Supplementary Figure 4) to the centroid for each condition. Abbreviations: EC = Escherichia coli, SD = Streptococcus dysgalactiae, SM = Streptococcus mitis, SP = Streptococcus parasanguinis, SPn = Streptococcus pneumoniae, SS = Streptococcus salivarius, STDDEV = standard deviation.

| **species** | **condition** | **number of PCs** | **variance explained [%]** | **mean distance** | **STDDEV distance** | **maximum distance** |
| --- | --- | --- | --- | --- | --- | --- |
| EC | Control | 2 | 88.3 | 0.68 | 0.33 | 1.02 |
| EC | IV10 | 2 | 88.3 | 3.50 | 1.52 | 5.25 |
| EC | MX10 | 2 | 88.3 | 0.24 | 0.16 | 0.36 |
| SD | Control | 2 | 98.9 | 0.12 | 0.09 | 0.17 |
| SD | IV10 | 2 | 98.9 | 0.21 | 0.14 | 0.31 |
| SD | MX10 | 2 | 98.9 | 0.43 | 0.19 | 0.64 |
| SM | Control | 2 | 98.9 | 0.12 | 0.06 | 0.18 |
| SM | IV10 | 2 | 98.9 | 0.25 | 0.11 | 0.37 |
| SM | MX10 | 2 | 98.9 | 0.10 | 0.03 | 0.13 |
| SP | Control | 2 | 99.2 | 0.08 | 0.03 | 0.12 |
| SP | IV10 | 2 | 99.2 | 0.11 | 0.05 | 0.16 |
| SP | MX10 | 2 | 99.2 | 0.14 | 0.07 | 0.19 |
| SPn | Control | 2 | 99.5 | 0.07 | 0.05 | 0.10 |
| SPn | IV10 | 2 | 99.5 | 0.34 | 0.15 | 0.51 |
| SPn | MX10 | 2 | 99.5 | 0.23 | 0.10 | 0.35 |
| SS | Control | 2 | 97.3 | 0.39 | 0.17 | 0.57 |
| SS | IV10 | 2 | 97.3 | 0.23 | 0.09 | 0.32 |
| SS | MX10 | 2 | 97.3 | 0.10 | 0.06 | 0.14 |

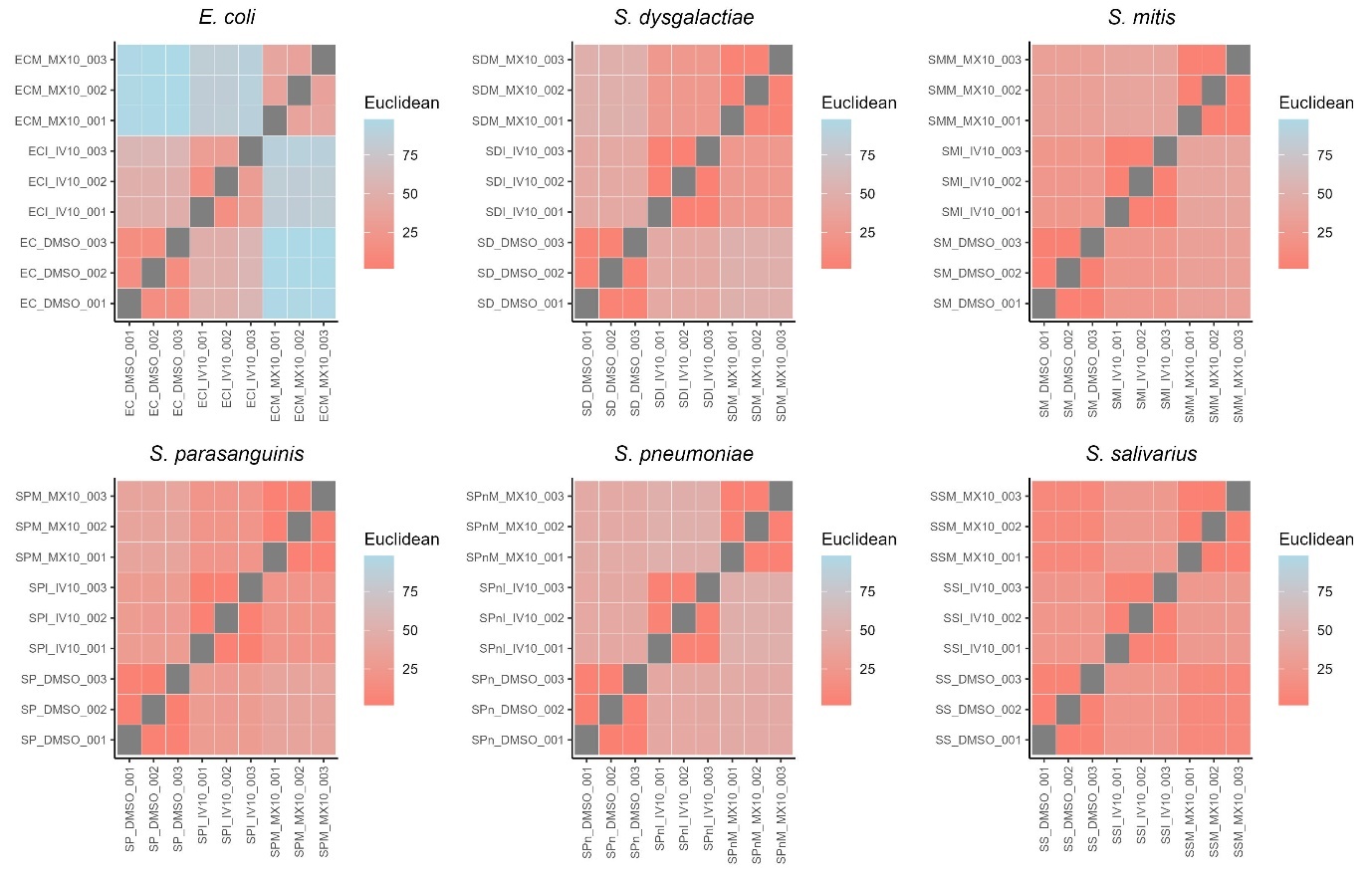

Supplementary Figure 5: Hierarchical clustering of 18 sample triplicates based on pairwise Euclidean distances calculated based on expression profiles.

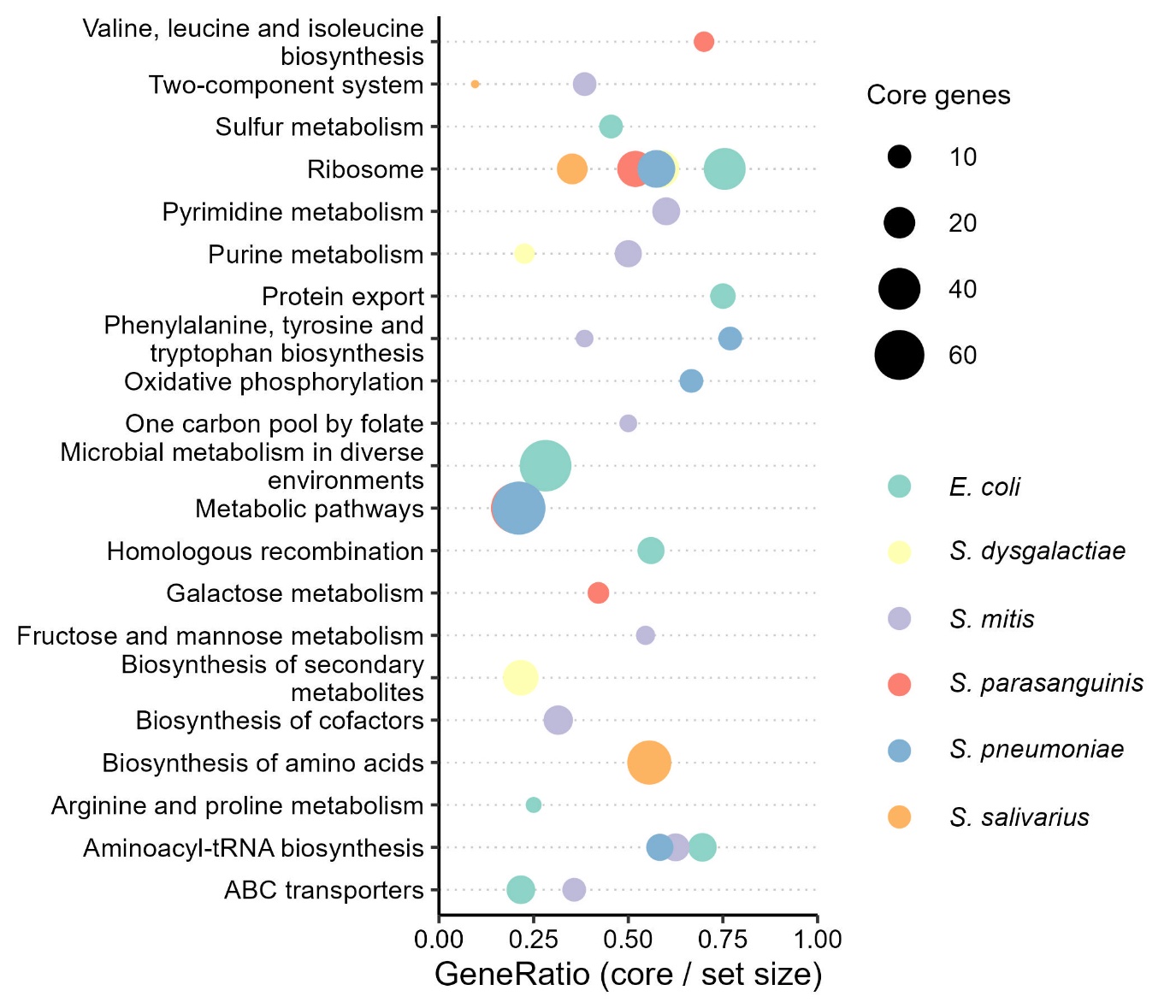

Supplementary Figure 6: Per-species dot plot of gene set enrichment analysis (GSEA). Gene-ratio is calculated as the “count of significantly enriched genes in a set / total genes in the set”.
